# Reduction of Spermine Synthase Suppresses Tau Accumulation Through Autophagy Modulation in Tauopathy

**DOI:** 10.1101/2023.03.17.533015

**Authors:** Xianzun Tao, Jiaqi Liu, Zoraida Diaz-Perez, Jackson R Foley, Tracy Murray Stewart, Robert A Casero, R. Grace Zhai

## Abstract

Tauopathy, including Alzheimer Disease (AD), is characterized by Tau protein accumulation and autophagy dysregulation. Emerging evidence connects polyamine metabolism with the autophagy pathway, however the role of polyamines in Tauopathy remains unclear. In the present study we investigated the role of spermine synthase (SMS) in autophagy regulation and tau protein processing in *Drosophila* and human cellular models of Tauopathy. Our previous study showed that *Drosophila spermine synthase* (*dSms*) deficiency impairs lysosomal function and blocks autophagy flux. Interestingly, partial loss-of-function of SMS in heterozygous *dSms* flies extends lifespan and improves the climbing performance of flies with human Tau (hTau) overexpression. Mechanistic analysis showed that heterozygous loss-of-function mutation of *dSms* reduces hTau protein accumulation through enhancing autophagic flux. Measurement of polyamine levels detected a mild elevation of spermidine in flies with heterozygous loss of *dSms*. SMS knock-down in human neuronal or glial cells also upregulates autophagic flux and reduces Tau protein accumulation. Proteomics analysis of postmortem brain tissue from AD patients showed a significant albeit modest elevation of SMS protein level in AD-relevant brain regions compared to that of control brains consistently across several datasets. Taken together, our study uncovers a correlation between SMS protein level and AD pathogenesis and reveals that SMS reduction upregulates autophagy, promotes Tau clearance, and reduces Tau protein accumulation. These findings provide a new potential therapeutic target of Tauopathy.

## INTRODUCTION

Tauopathy is a group of neurological disorders, including Alzheimer Disease (AD), Progressive Supranuclear Palsy, Chronic Traumatic Encephalopathy and so on, characterized by abnormal Tau accumulation in the brain [1–3]. Accumulated Tau species cause neuronal damage and death [4, 5]. Protein degradation machineries, including proteasome and autophagy, play important roles in Tau homeostasis [6–8]. Abnormal accumulation of intermediate autophagy structures is observed in brains of AD patients or animal models, suggesting autophagic flux is blocked in AD [9–12]. Reducing neurotoxic Tau accumulation by repairing or enhancing autophagy is of strong potential to treat AD or other Tauopathies.

Recent studies have indicated the potential role of polyamines in autophagy and Tauopathy [13–17]. Polyamines are positively charged alkylamines, including spermidine and spermine, and their precursor putrescine. Through interacting with negatively charged DNA, RNA or proteins, they are broadly involved in cellular activities. Polyamines can be accumulated from the extracellular environments or synthesized in the cells [18, 19]. Mutations of polyamine transporters or metabolic enzymes are related to diseases [20–23]. For example, mutations in *spermine synthase (SMS)* cause Snyder-Robinson Syndrome (SRS), characterized by developmental delay, muscle/bone abnormalities, and intellectual disability [24]. Mechanistic studies showed that SMS deficiency results in accumulation of spermidine and its metabolites [21, 25]. Increased byproducts of spermidine catabolism, aldehydes and ROS, damage membrane structures in the cells, such as mitochondria and lysosomes [21, 26].

Polyamine metabolism appears to be linked to the intersection of Tauopathy and autophagy. For example, polyamine levels and the polyamine metabolic pathway are upregulated in brains of Tauopathy patients or animal models [13, 14, 27, 28]. Manipulating polyamine metabolic enzymes has been shown to modulate Tauopathy progression in animal models [13, 14]. The regulation of autophagy by polyamines is complex. On one hand, increasing cellular polyamine levels by spermidine/spermine supplementation or overexpression of putrescine synthase ornithine decarboxylase 1 (ODC1) enhances autophagy through upregulating autophagy-related genes [15, 29]. On the other hand, polyamine imbalance and spermidine accumulation in SMS-deficient cells lead to lysosomal damage and blocked autophagic flux [21]. The underlying mechanism of these seemingly contradictory effects of polyamines on autophagy remains unclear, which hinders the potential of targeting the polyamine pathway to treat highly autophagy-related diseases, such as Tauopathy.

In the present study, we explored the interaction between SMS and autophagy under Tauopathy conditions. We revealed distinctive effects of heterozygous or homozygous loss-of-function *dSms* mutations in *Drosophila* and an unexpected protective effect of SMS reduction against Tauopathy in a *Drosophila* model and human cells. We further examined expression-level alterations of polyamine metabolism enzymes in AD brains and propose SMS as a potential therapeutical target of Tauopathy.

## RESULTS

### SMS reduction ameliorates Tauopathy in a *Drosophila* model

The impact of polyamine catabolism on autophagy/lysosomal flux suggests that the polyamine metabolic pathway and protein homeostasis are closely connected [21, 26]. Given the importance of protein homeostasis in neurodegenerative Tauopathy, this connection suggests that polyamine metabolism might play a role in modulating disease progression in Tauopathy, where dysregulated protein homeostasis and autophagic flux are pathological hallmarks [9–12]. To test the effect of SMS reduction in Tauopathy, we established a *Drosophila* line with human Tau (hTau) overexpression and a heterozygous loss-of-function mutation of *dSms (dSms^+/-^*) [4, 21]. As reported previously, compared to control lacZ-overexpressing flies, hTau-overexpressing flies showed significantly reduced lifespan and impaired locomotor behavior [4] (Figure 1A, 1B and S1A and S1B). Interestingly, introducing the heterozygous *dSms*^+/-^ mutation significantly extended the lifespan of hTau-expressing flies (Figure 1A and S1A) and improved the age-dependent behavior impairment (Figure 1B and S1B).

**Figure 1.**
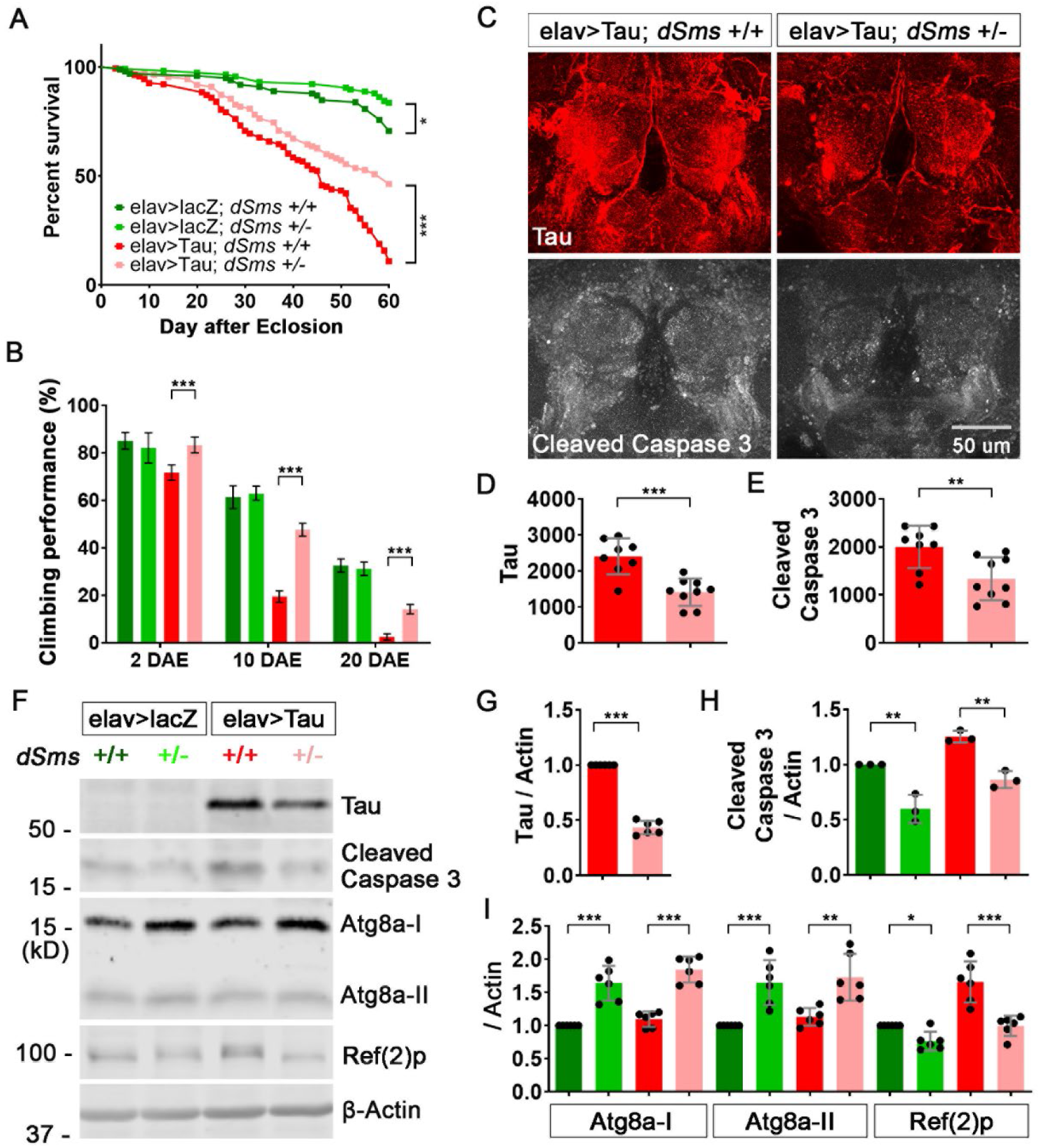
SMS reduction ameliorates Tauopathy in a *Drosophila* model. (A) Lifespan of female flies with indicated genotype. n = 99, 116, 164, 110; Log-rank (Mantel-Cox) test. (B) Climbing performance of female flies with indicated genotype at indicated ages. n = 100, 100, 100, 100. (C) Staining of brains of 10 DAE flies with antibodies against Tau or cleaved caspase 3. The image is a representative of multiple brains in each group, n = 8, 9. (D, E) Quantification of the staining signal intensity of Tau (D) or cleaved caspase 3 (E) in (C). n = 8, 9. (F) Western-blot of Tau, cleaved caspase 3, autophagy marker Atg8a and Ref(2)p in heads of 10 DAE flies. The image is a representative of multiple experiments. (G) Quantification of the protein level of Tau (G), cleaved caspase 3 (H) or autophagy markers (I) in (F). All the protein levels were normalized with the β-Actin level. All the values were further normalized by that of the control flies. n = 6, 3, 6. (B, H, I) Two-way ANOVA multiple comparisons. (D, E, G) Student’s *t* test. Data represent mean ± SEM.

To dissect the cellular and molecular mechanisms underlying the beneficial effects of heterozygous loss of *dSms*, we first evaluated the level of hTau protein accumulation in fly brains. Staining the brains with antibodies against hTau proteins showed that hTau proteins were significantly reduced in the fly brains with heterozygous *dSms*^+/-^ (Figure 1C and 1D). Furthermore, the downstream neuronal toxicity effector of Tauopathy, cleaved caspase 3, significantly decreased in the brains with heterozygous *dSms^+/-^* (Figure 1C and 1E). Western-blot analysis confirmed the decrease of hTau and cleaved caspase 3 in the heads of flies with heterozygous *dSms^+/-^* (Figure 1F-H).

Since Tau protein homeostasis is regulated by autophagy [30–32], we wondered whether autophagic flux is altered with heterozygous loss of *dSms*. We used several different autophagy markers to evaluate the level of autophagic flux, including Atg8a-I and its lipidated form, Atg8a-II, which are cytoplasmic and autophagosome-associated, respectively [33, 34]; and Ref(2)p, the cargo recruiter of autophagy [35]. As shown in Figure 1F and 1I, Atg8a-I and its lipidated form, Atg8a-II were significantly upregulated in heterozygous *dSms* fly brains. Interestingly, Ref(2)p was significantly downregulated in the brains of heterozygous *dSms* flies with either lacZ or hTau overexpression (Figure 1F and 1I), suggesting that heterozygous loss of *dSms* enhances autophagic flux independent of hTau overexpression. This enhanced autophagic flux with partial loss of dSms is likely the underlying mechanism of the lifespan-extending effect of heterozygous loss of *dSms* in either lacZ or hTau-overexpressing flies (Figure 1A and S1A). Collectively, these results show that partial loss of SMS enhances autophagic flux, extends lifespan, and ameliorates neurodegenerative phenotypes in Tauopathy models.

### SMS regulates autophagy in a biphasic manner

The neuroprotective effects observed in these dSMS heterozygotes are in contrast to our previous observations in dSMS homozygous mutants, where complete loss of dSMS (*dSms^-/-^*) resulted in elevated spermidine level and impaired autophagic flux [21]. To exclude the possible interference by lacZ or hTau overexpression, we compared the autophagy level in flies with heterozygous or homozygous *dSms* without lacZ or hTau overexpression (genotype: heterozygous *dSms*^+/-^, or homozygous *dSms*^-/-^). Strikingly, autophagic flux alteration in heterozygous or homozygous flies occurred in opposite directions: while Atg8a-I (cytoplasmic) was increased in either homozygous or heterozygous flies, Atg8a-II (autophagosome-associated) was reduced in homozygous flies but increased in heterozygous flies (Figure 2A and 2B). In addition, the autophagy cargo recruiter Ref(2)p was accumulated in homozygous flies but reduced in heterozygous flies (Figure 2A and 2B). Taken together, these data suggest that autophagic flux is blocked in homozygous flies, as reported in our previous study, but is upregulated in heterozygous flies.

**Figure 2.**
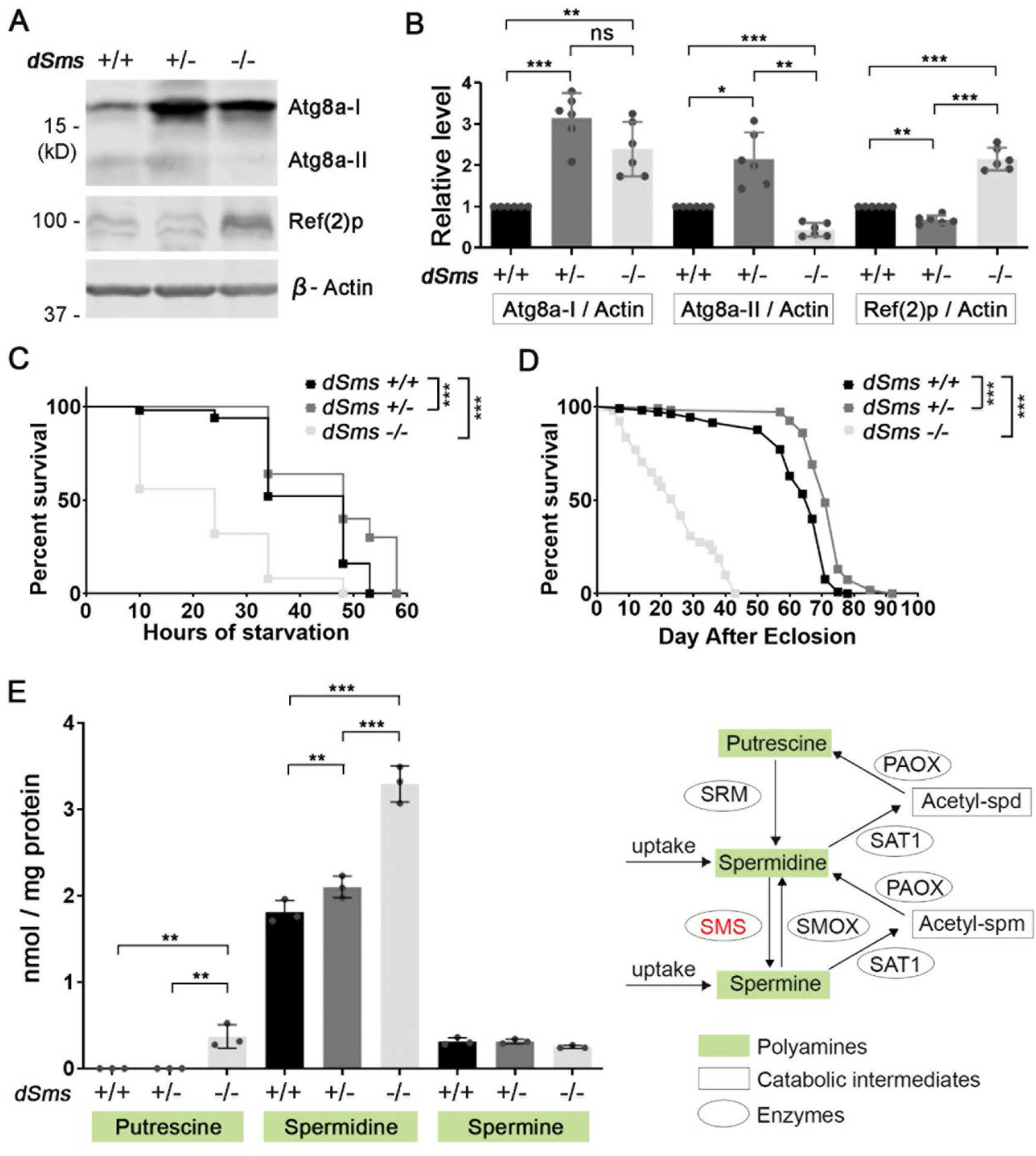
SMS regulates autophagy in a bi-phasic manner. (A) Western-blot of autophagy marker Atg8a and cargo recruiter Ref(2)p in 5 DAE flies. The image is a representative of multiple separate experiments. (B) Quantification of the protein level of Atg8a-I, Atg8a-II or Ref(2)p in (A). All the protein levels were normalized with the β-Actin level. All the values were further normalized by that of the control flies. n = 6; two-way ANOVA multiple comparisons. (C) Survival curve of 10 DAE female flies with indicated genotype under starvation. n = 50, 50, 25; Log-rank (Mantel-Cox) test. (D) Lifespan of female flies with indicated genotype. n = 105, 107, 91; Log-rank (Mantel-Cox) test. (E) Polyamine levels of 10 DAE female flies with indicated genotype. Each dot indicates a sample of homogenized mixture of 10 flies. n = 3; two-way ANOVA multiple comparisons. Data represent mean ± SEM. The measurement showed here was done together with that showed in our previous publication [26]. The data of the control and *dSms* -/- flies are shared in these two studies.

To further assess the functional consequence of autophagy alteration, we evaluated the starvation resistance of these flies, which is highly dependent on autophagy activity. While homozygous flies died earlier under starvation compared to control flies, heterozygous flies survived longer than control flies (Figure 2C and S2A). We then determined the lifespan of these flies, which is also related to autophagy activity. While homozygous flies lived much shorter than control flies, heterozygous flies lived significantly longer than control flies (Figure 2D and S2B).

We next questioned whether the different autophagic activity alteration in heterozygous and homozygous flies results from different polyamine level alteration in these flies. To test the possibility, we measured the polyamine levels in these flies. Interestingly, the level of spermidine is elevated in either homozygous or heterozygous flies compared to that in the control flies (Figure 2E and S2C). But the spermidine level increase in heterozygous flies is much milder than that in homozygous flies (Figure 2E and S2C). The spermine level is reduced in homozygous flies, as expected, but not altered in heterozygous flies (Figure 2E and S2C). Collectively, these data suggest that SMS regulates autophagy through modulating spermidine. Specifically, high level of spermidine accumulation by homozygous deletion of *dSms* impairs autophagic flux but mild spermidine elevation by heterozygous deletion of *dSms* upregulates autophagic flux.

### SMS reduction upregulates autophagic flux in both neurons and glia in *Drosophila*

To further validate the regulation of SMS on autophagic flux in single cells in vivo, we used an Atg8a reporter tagged with a pH-insensitive mCherry and a pH-sensitive GFP [36] to monitor autophagy flux (Figure 3A). Before fusion with acidic lysosomes, both mCherry and GFP fluorescence from the reporter proteins, in cytosol or on the autophagosome membrane, are stable. However, after fusion with acidic lysosomes, the pH decrease causes a reduction in GFP fluorescence. As autolysosomes mature, the reporter proteins start to be degraded, accompanied by mCherry fluorescence reduction (Figure 3A). As such, the intensity and the ratio of the two fluorescence signals can be used to differentiate autophagic structures. In phagophores or autophagosomes, both mCherry and GFP signals are high and the ratio of mCherry to GFP is low. In newly formed autolysosomes, the mCherry signal is high, the GFP signal is low and the ratio is high. In matured autolysosomes, the mCherry signal is medium, the GFP signal is low and the ratio is medium. By measuring the ratio of total mCherry fluorescence to total GFP fluorescence in a single cell, we can roughly evaluate the autophagic flux in the cell: higher ratio indicates more functional autolysosome enrichment (Figure 3B). Based on the fluorescence intensity of mCherry and GFP in a scatter plot, cells can be divided into three populations: high-mCherry/high-GFP: cells enriched with phagophores/autophagosomes or free reporter proteins; high-mCherry/low-GFP: cells enriched with functional autolysosomes; and low-mCherry/low-GFP, cells with matured autolysosome enriched or low expression of the reporter proteins (Figure 3C).

**Figure 3.**
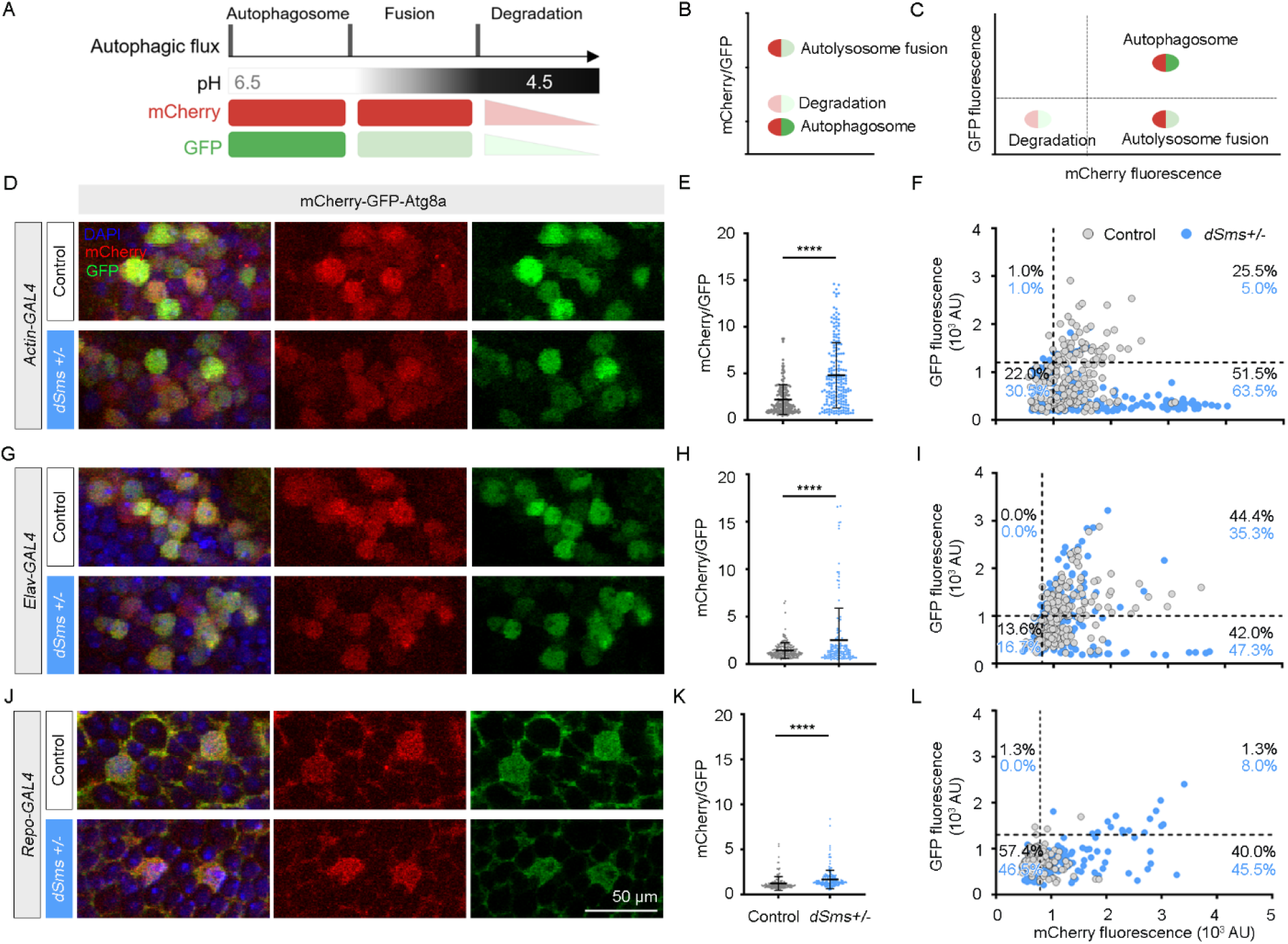
SMS reduction upregulates autophagic flux in both neurons and glia in *Drosophila*. (A) Diagram of the fluorescence signals from the reporter protein mCherry-GFP-Atg8a under different conditions. (B) The ratio of total fluorescence intensity of mCherry to GFP in single cells indicates the enrichment of specific stage of autophagic flux, which can be used to evaluate the average autophagic flux level in a cell. (C) Scat plotting of mCherry and GFP fluorescence intensity divides cells into subgroups enriched with specific stage of autophagic flux. (D, G, J) Images of brain cells of control or *dSms^+/-^* flies with *actin-, elav-* or *repo-driven* mCherry-GFP-Atg8a expression, 10 DAE, female flies. (E, H, K) The ratio of total fluorescence intensity of mCherry to GFP in single cells in the brains from (D), (G) and (J), respectively. (F, I, L) Grouping cells with the florescence ratio in single cells from (D), (G) and (J), respectively. Each dot represents a single cell. 50 cells from each brain, 3 or 4 brains from each group, were measured. n = 150 or 200; Student’s *t* test. Data represent mean ± SEM.

To uncover potential cell type-specific alterations in autophagy in the brain, we established *Drosophila* lines expressing the mCherry-GFP-Atg8a reporter controlled by different drivers (*actin-GAL4, for global expression, elav-GAL4 for neuronal specific expression* and *repo-GAL4 for glial expression)*, with or without partial loss of *dSms* (Figure 3D, 3D, and 3J). Interestingly, we found that regardless of driver control, the ratio of mCherry to GFP fluorescence in the brain cells of the flies with heterozygous loss of *dSms* is significantly higher than that of the control flies (Figure 3E, 3H, and 3K), suggesting SMS reduction upregulates autophagic flux in both neurons and glia. Notably, the general intensity of mCherry or GFP signals in brain cells of heterozygous flies is lower than that of control flies (Figure 3D, 3G, and 3J), which indicates the potentially higher autophagic flux in the heterozygous flies, although we couldn’t exclude the possibility that the expression level of the reporter protein is somehow lower in the heterozygous flies. Cell populations of high-mCherry/low-GFP in *dSms^+/-^* heterozygous flies are significantly larger than that in control flies in all the three cell type-specific expressing conditions (Figure 3F, 3I, and 3L), suggesting SMS reduction increases functional autolysosomes in all cells.

Notably, we found that compared to the neuronal population (driven by elav-GAL4), glial cells (driven by repo-GAL4) have significantly larger low-mCherry/low-GFP cell population but smaller high-mCherry/high-GFP cell populations (Figure 3I and 3L). This might indicate a higher rate of matured autolysosomes in glia and/or less accumulated reporter proteins because of a shorter lifespan of glia. Taken together, SMS reduction upregulates autophagic flux in both neurons and glia, even though the autophagic flux itself might be different in the two types of cells.

### SMS knock-down reduces Tau accumulation in human neuronal or glial cell lines

We then tested the effect of SMS reduction in human neuron-like cells. SMS knock-down with siRNA in SH-SY5Y cells mildly upregulated LC3-I and LC3-II (the human homologues of *Drosophila* Atg8a-I and Atg8a-II), downregulated p62 (the human homologue of *Drosophila* Ref(2)p [35]) and significantly decreased exogenous Tau protein accumulation (Figure 4A and 4B). Notably, overexpressed EGFP was also significantly downregulated by SMS knock-down (Figure 4A and 4B), suggesting the effect of SMS knock-down on autophagy is not Tau-specific but rather a general effect on cellular protein homeostasis. Given that glial cells have been shown to play important roles in Tau aggregate regulation [37, 38], we next investigated the effect of SMS reduction on Tau aggregates in human glial cells using a Tau K18 fibrils uptake assay [32]. We incubated control or SMS siRNA-knockdown SVG p12 cells with Tau K18 fibrils conjugated with Alexa-fluor-647. Consistent with previous report [32], Tau K18 fibrils were taken up by SVG p12 cells efficiently (Figure 4C). In addition, we observed colocalization of Tau K18 fibrils and p62 (Figure 4C), suggesting the regulation of Tau fibrils by autophagy. As shown in Figure 4C-D, SMS knock-down significantly decreased the size and the intensity of Tau fibril loci in SVG p12 cells. This result is consistent with the observation of reduction of Tau accumulation with partial loss of SMS in both human neuronal SH-SY5Y cells and *Drosophila* in vivo models.

**Figure 4.**
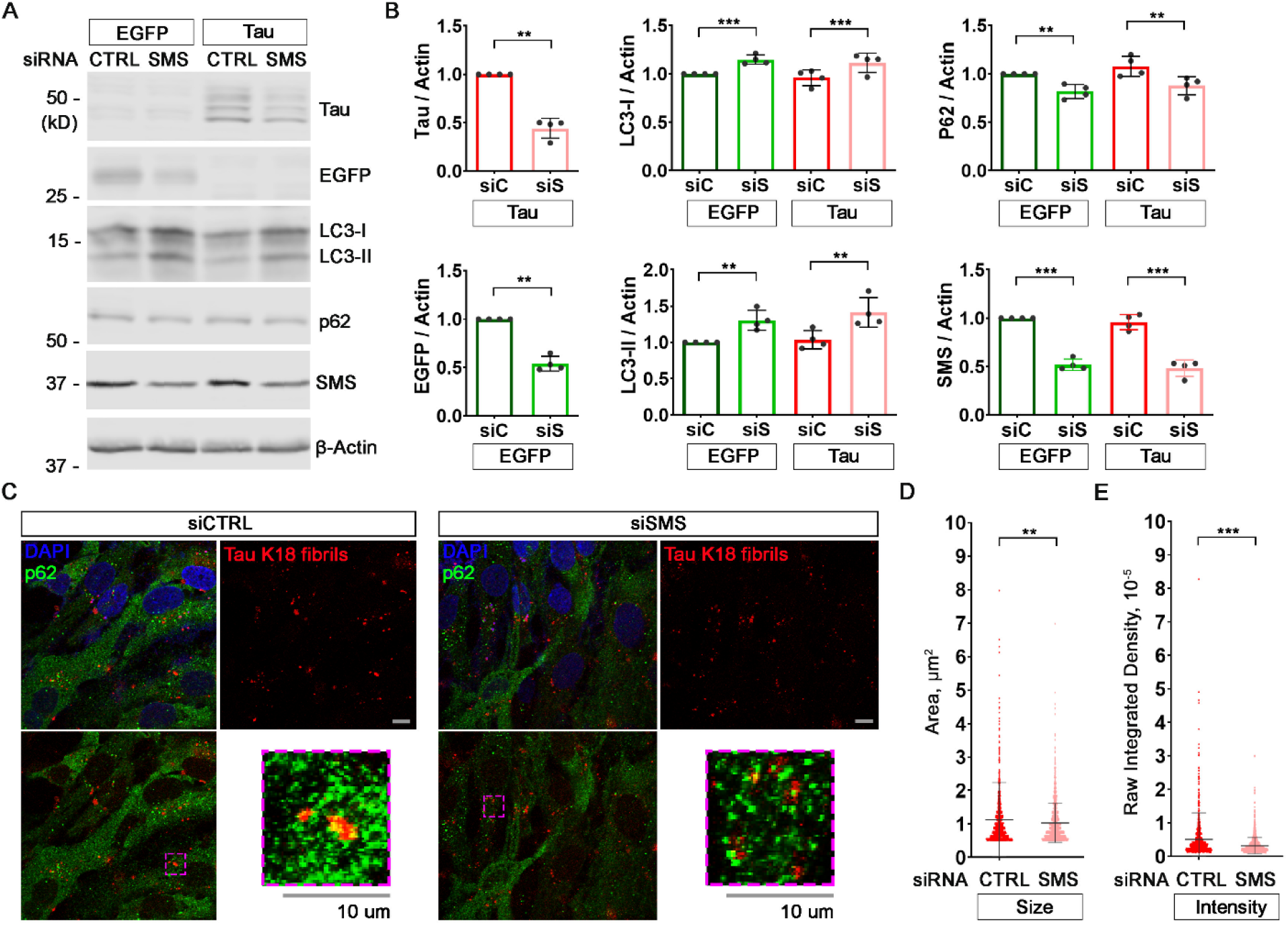
SMS knock-down upregulates autophagy and suppresses Tau accumulation in human neuronal or glial cell lines. (A) Western-blot of Tau, EGFP, autophagy marker LC3, cargo recruiter p62 and SMS in SH-SY5Y cells with Tau/EGFP plasmids and Control/SMS siRNA transfection. The image is a representative of four separate experiments. (B) Quantification of the protein levels of Tau (5A6), EGFP, LC3-I (cytoplasmic), LC3-II (autophagosome-associated), p62 and SMS in (A). All the protein levels were normalized with the β-Actin level. All the values were further normalized by that of the control cells. n = 4; Student’s *t* test (Tau or EGFP) or two-way ANOVA multiple comparisons (others). (C) p62 staining and Alexa 647-conjugated Tau K18 fibrils in SVG p12 cells with Control/SMS siRNA transfection. The images are representatives of five fields. (D) Quantification of the size (area) and intensity of the Alexa 647 conjugated Tau K18 fibrils in (C). n = 5; Student’s t test. Data represent mean ± SEM.

### SMS levels are elevated in post-mortem AD patient brains

To directly assess the potential correlation of SMS and polyamine metabolism with AD, we examined the expression level of SMS and other polyamine pathway enzymes (Figure 5A) in the frontal cortex of AD patient brains using a meta-analysis [39] of seven published proteomic datasets based on the TMT-LC/LC-MS/MS platform [40–43]. The protein level of SMS is consistently upregulated in trend in AD brains in all seven datasets, and the combined p value analysis shows the upregulation is significant (Figure 5B). In contrast, the trend of the change of the protein level of spermidine synthase (SRM) in AD brains varies among the datasets, and the combined analysis suggests it is downregulated in AD brains (Figure 5C). The protein levels of the polyamine catabolic enzymes are under the detection limit in some datasets and the alterations vary among the datasets (Figure 5D, 5E and 5F). However, combined analysis shows that both spermine oxidase (SMOX) and spermidine/spermine acetyltransferase (SAT1) are upregulated in AD brains. Collectively, it suggests that spermidine and spermine interconversions mediated by SMS and spermine catabolic enzymes are elevated in AD, which likely result in the accumulation of potentially toxic metabolites and increased oxidative stress in the brain.

**Figure 5.**
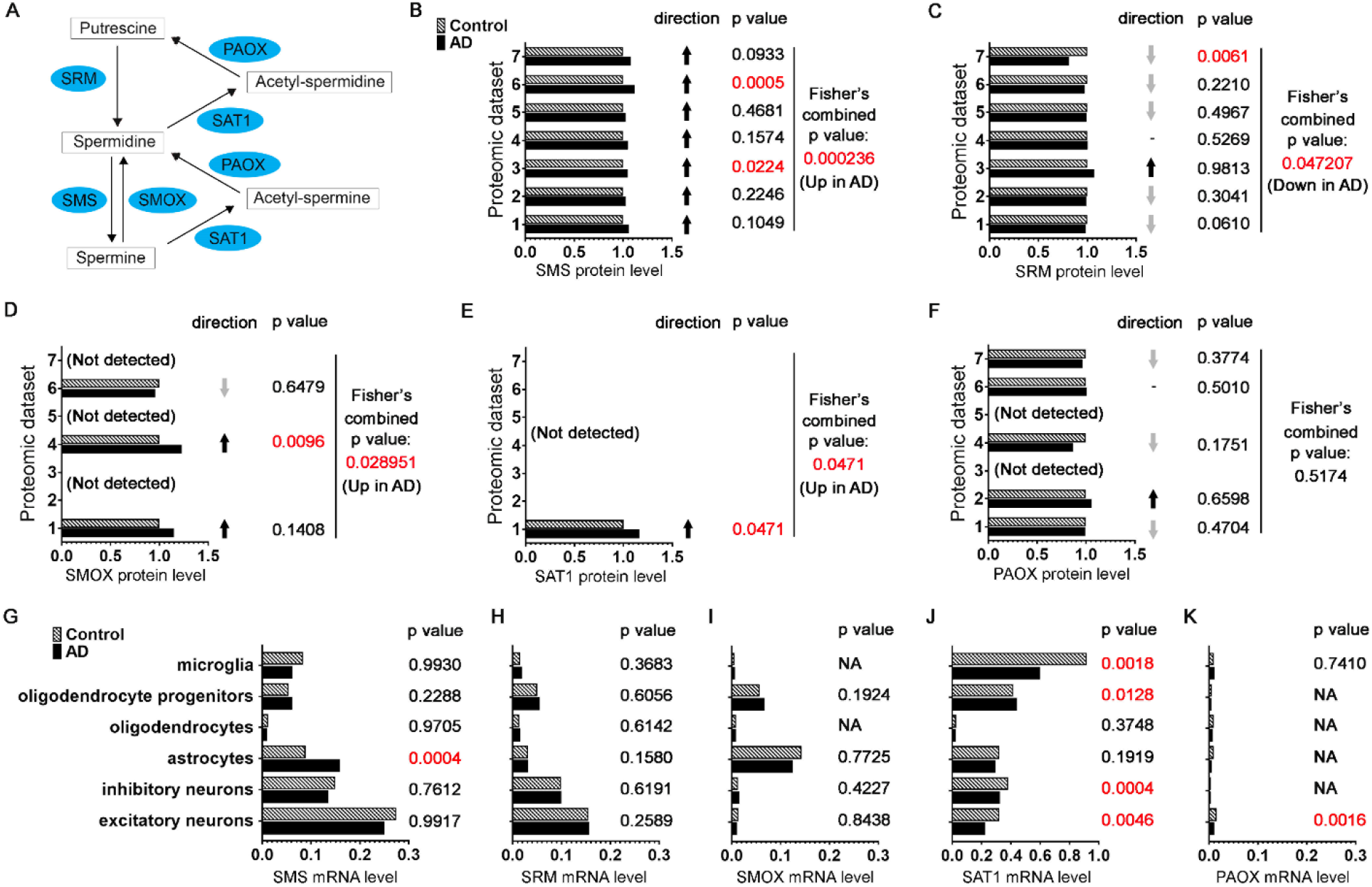
SMS expression level is upregulated in post-mortem AD patient brains. (A) Diagram of polyamine pathway with the enzymes highlighted. (B-F) The protein levels of SMS (B), SRM (C), SMOX (D), SAT1 (E) or PAOX (F) in frontal cortex of control or AD patient brains measured by quantitative mass spectrometry from seven published datasets [39–43]. The one-tailed *t*-test p-values was combined with Fisher’s method. (G-K) The mRNA levels of SMS (G), SRM (H), SMOX (I), SAT1 (J) or PAOX (K) in indicated brain cells from prefrontal cortex of control or AD patients measured by single-nucleus RNA seq reported recently [44]. The p values were obtained with the Poisson mixed model.

To further uncover potential cell type-specific changes of the expression level of SMS and other polyamine pathway enzymes in the brain, we analyzed a published single-nucleus RNA seq dataset from prefrontal cortex samples of control or AD patients [44]. Interestingly, the mRNA level of SMS is significantly upregulated in astrocytes of AD patients but is largely unchanged in other cell types (Figure 5G). As a comparison, the mRNA level of SRM is not significantly altered in all the measured cell types (Figure 5H). The mRNA levels of polyamine catabolic enzymes are not significantly changed in astrocytes of AD patients (Figure 5I, 5J and 5K), further highlighting the specificity of SMS mRNA level upregulation in AD. It would be interesting to detect the protein levels of polyamine pathway enzymes in specific cell types in AD in future studies.

## DISCUSSION

In the present study, we investigated the regulation of SMS on autophagy in physiological or disease conditions (Figure 1, 2 and 3). We found that heterozygous loss-of-function of *dSms* significantly upregulates autophagy, decreases hTau protein accumulation and ameliorates Tauopathy in a *Drosophila* model (Figure 1). Consistently, SMS knock-down in human neuronal or glial cells also enhances autophagic activity and decreases exogenous Tau accumulation (Figure 4). Moreover, we showed that SMS, together with some polyamine catabolic enzymes, is elevated in AD patient brains (Figure 5), suggesting SMS plays roles in AD and could be a therapeutic target.

Our previous studies in flies with homozygous loss-of-function mutation of *dSms* modeling Snyder-Robinson syndrome discovered the detrimental effect of accumulated spermidine on autophagic flux, caused by increased aldehyde-mediated lysosomal damage [21, 26]. This observation is against the beneficial effect of polyamine supplementation on autophagy tested in multiple organisms [15, 16]. In the present study, we found opposing effects of partial versus complete loss of *dSms* on autophagic activity (Figure 2), which provides a possible explanation to the “controversy” between accumulated spermidine in Snyder-Robson syndrome and that from polyamine supplementation. It is likely that increased spermidine enhances autophagy gene expression in both conditions (Figure 2A and 2B). However, the higher level of spermidine accumulation results in abnormal accumulation of the catabolic byproducts, including aldehydes and ROS, and therefore causes lysosomal damage and the subsequent block of autophagic flux in Snyder-Robinson syndrome (Figure 6).

**Figure 6.**
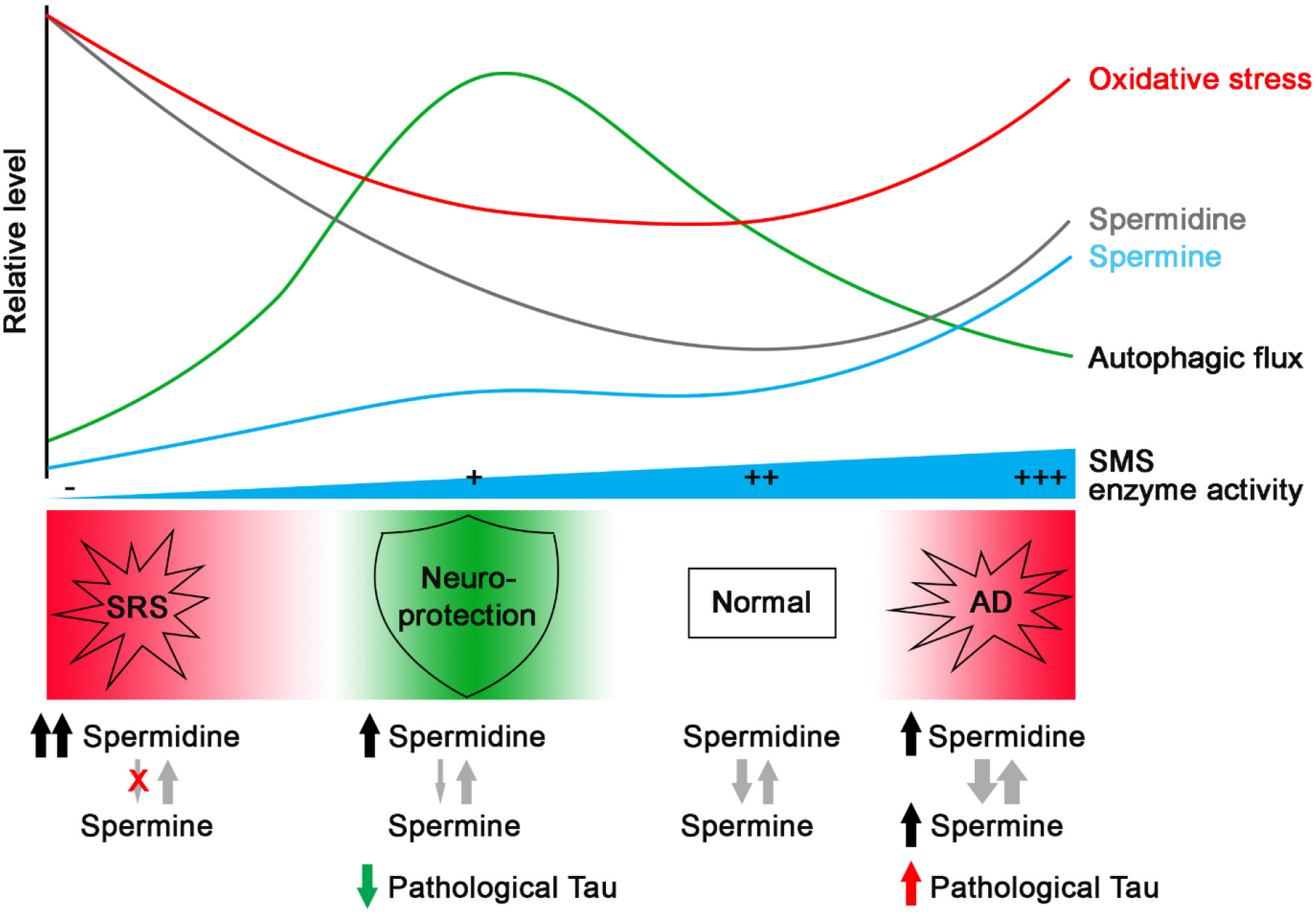
Model of interaction of polyamine pathway and autophagy in SRS and Tauopathy. Under SRS conditions, with SMS deficiency, there is an increase in spermidine and a decrease in spermine levels, which leads to an overall increase in total polyamines. Conversely, under AD conditions, with increased SMS and the conversion of spermine to spermidine, both spermidine and spermine levels increase. The elevated polyamines and their metabolism in these two pathological conditions result in higher levels of oxidative stress and impaired autophagic flux. However, a partial reduction of SMS leads to a mild accumulation of spermidine without significant changes in spermine levels, which avoids triggering oxidative stress and enhances autophagic flux. While impaired autophagic flux in AD aggravates pathological Tau accumulation, enhancing autophagic flux through SMS partial reduction promotes Tau clearance and confers neuroprotection.

The observation of elevated polyamine levels in AD patient brains or plasma [27, 28, 45–47] also raises questions on the perceived beneficial effect of spermidine supplementation on brain health in human or animal models [29, 48–51]. Different levels of polyamine metabolic activity might contribute to this discrepancy. Consistent with the upregulation of polyamine metabolism enzymes in AD patient brains (Figure 4), the polyamine pathway in Tauopathy animal models is activated [13, 14]. It is likely that high levels of polyamine catabolism, such as SMOX-mediated spermine back-conversion to spermidine, results in oxidative stress in Tauopathy, whose detrimental effect overcomes the beneficial effect of polyamine themselves (Figure 6). An appropriate level of exogenous polyamine supplementation might be able to assert the beneficial effect of polyamines without triggering the activation of the polyamine catabolic pathway, due to moderate levels of cellular accumulation. It would be interesting to test the effects of the combination of polyamine supplementation and inhibition of polyamine catabolism on Tauopathy.

In summary, we have demonstrated that SMS regulates autophagic activity in a complex manner, probably through modulating the level of polyamines and the catabolic process. While severe to complete loss or overexpression of SMS causes significant polyamine dysregulation and autophagy blockade, partial reduction of SMS leads to mildly accumulated spermidine and enhanced autophagic activity, promotes Tau clearance and confers neuroprotection. Our discovery suggests SMS as a potential therapeutical target of Tauopathy such as AD.

## METHODS

### *Drosophila* stocks and genetics

Unless specified, flies were maintained on a cornmeal-molasses-yeast medium at 25 °C, 65% humidity, 12 h light/12 h dark. The following fly strains were used in the studies: yw, actin-GAL4, elav-GAL4, repo-GAL4, USA-lacZ, USA-Tau, UAS-mCherry-GFP-Atg8a (Bloomington Drosophila Stock Center); CG4300^e00382^ (The Exelixis Collection, Harvard Medical School).

### Fly lifespan assay

Newly eclosed flies were collected and about 20 flies of the same sex were kept in a fresh food vial. Flies were transferred to new vials every week and the number of live flies was counted every other day.

### Fly climbing assay

Age-matched female or male flies from each genotype were placed in a vial marked with a line 8 cm from the bottom surface. The flies were gently tapped onto the bottom and given 10 s to climb. After 10 s, the number of flies that successfully climbed above the 8 cm mark was recorded and divided by the total number of flies. The assay was repeated 10 times, and 10 independent groups (total 100 flies) from each genotype were tested.

### Fly starvation resistance assay

Age-matched female or male flies were placed in a vial with three pieces of filter paper soaked with water; the filter paper was kept wet the entire process. The number of dead flies was counted every two hours.

### Brain dissection and immunohistochemical staining

Flies were dissected in cold PBS (pH =7.4). Brains were fixed in freshly made 4% formaldehyde for 15 min. After 10 min washing in PBS containing 0.4% (v/v) Triton X-100 (PBTX) for three times, brains were incubated with primary antibodies diluted in 0.4% PBTX containing 5% goat serum on a roller at 4 °C overnight. Brains were washed in 0.4% PBTX 10 min for three times and then incubated with conjugated secondary antibodies on a roller at 4 °C overnight, followed by 10 min washing in 0.4% PBTX and then staining with DAPI for 10 min. After 10 min washing in 0.4% PBTX, brains were mounted on glass slides with VECTASHIELD Antifade Mounting Medium (Vector Laboratories) and kept at 4 °C until imaging. Brains were imaged using an Olympus IX81 confocal microscope coupled with ×10, ×20 air lens or ×40, ×60 oil immersion objectives. Images were processed using FluoView 10-ASW software (Olympus) and analyzed using ImageJ software.

### Immunoblot analysis

For immunoblot analysis of proteins, tissues were homogenized in RIPA buffer (R0278, Thermo) with proteinase inhibitors (11836170001, Roche) and phosphatase inhibitors (04906837001, Roche) on ice, followed by 10 min centrifUgation at 10,000×g at 4 °C. The supernatants were then mixed with Laemmli sample buffer and heated at 95 °C for 10 min. Proteins were separated on a Bis-Tris gel and transferred to a nitrocellulose membrane. After blocking, the membrane was incubated with primary antibodies overnight at 4 °C and near infrared dye-conjugated secondary antibodies for 2 h at room temperature. Imaging was carried out on an Odyssey Infrared Imaging system (LI-COR Biosciences) and images were analyzed using Image Studio software.

### Antibodies

The following commercially available antibodies were used: anti-GABARAP for Drosophila Atg8a (PM037, MBL), anti-Ref(2)P (ab178440, Abcam), anti-Tau 5A6 (5A6, DSHB), anti-Phospho-Tau AT8 (MN1020, Thermo), anti-cleaved caspase 3 (9661, Cell Signaling), anti-LC3B (L7543, Sigma), anti-p62 (NBP1-48320, Novus Biologicals), anti-GFP (G5144, Invitrogen), anti-Actin (A1978, Sigma), Cy5-conjugated anti-HRP (123175021, Jackson ImmunoLab), and secondary antibodies conjugated to Alexa 488/568/647 (Thermo Fisher Scientific), or near infrared (IR) dye 700/800 (Rockland).

### Polyamine extraction and measurements by HPLC

Samples were collected from flash-frozen flies stored at −80 °C. Polyamine content was determined by the pre-column dansylation, high-performance liquid chromatography method of Kabra, et al. using 1,7 diaminoheptane as the internal standard [52]. The measurement shown here was done together with that shown in our previous publication [26]. The data of the control and *dSms* -/- flies are shared in these two studies.

### SMS knock-down in human cells

Co-transfections of siRNA (s13173, ThermoFisher) and plasmids expressing EGFP (pEGFP-C1, Clontech) or Tau into SH-SY5Y cells (CRL-2266, ATCC) was performed with JetPRIME transfection reagent (114-07, Polyplus) according to the manufacturer’s instruction in a 12-well plate. After 24 hours, replace the medium, followed by another medium change after 48 hours. On the fifth day, harvest the cells with RIPA buffer (R0278, Thermo) with proteinase inhibitors (11836170001, Roche) and phosphatase inhibitors (04906837001, Roche) on ice. The plasmid expressing Tau was subcloned from pRK5-EGFP-Tau (46904, Addgene) with the Tau coding sequence replacing the EGFP-Tau fusion sequence between the restriction sites ClaI and SalI.

### Tau K18 fibril labeling and transformation

Tau K18 fibrils (NBP2-76793, NovusBio) were labeled with Alexa Fluor succinimidyl ester dye 647 (A37573, Thermo) and transformed into cells as reported [32]. Control or SMS siRNA are transfected into SVG p12 cells (CRL-8621, ATCC) on coverslips in 12-well plates as mentioned above. Three days after transfection, the sonicated Tau K18 fibrils were added into the medium. Two days later, the cells were fixed with 4% FA for 15 minutes, permeabilized with 0.4% Triton X-100 for 5 minutes and stained with p62 antibodies and DAPI.

### Statistics

Data were analyzed with Prism (GraphPad Software). Log-rank (Mantel-Cox) test with correction for multiple comparisons (Bonferroni method) was used for survival curve (lifespan) analysis. Student’s *t* test (two-tailed) was used for comparison of two groups of samples for other assays. One-way ANOVA multiple comparisons was used for assays with more than two groups. A p value smaller than 0.05 is considered statistically significant. * indicates p < 0.05. ** indicates p < 0.01. *** indicates p < 0.001.

## Funding

National Institutes of Health grant R01NS109640 to RGZ.

National Institutes of Health grants R01HD110500 to RAC and TMS and R01CA235862 to RAC

University of Pennsylvania Orphan Disease Center Million Dollar Bike Ride grants MDBR-20-

135-SRS and MDBR-21-106-SRS to RAC and TMS

The Chan Zuckerberg Initiative (to RAC and TMS)

## Author contributions

Conceptualization: XT, RGZ

Methodology: XT, JL, ZDP, JRF, TMS, RACJ, RGZ

Investigation: XT, JL, ZDP, JRF, TMS, RACJ, RGZ

Visualization: XT, RGZ

Funding acquisition: RGZ

Project administration: RGZ

Supervision: RGZ

Writing – original draft: XT, RGZ

Writing – review & editing: YZ, TMS, RACJ, HW

## Data and materials availability

All data are available in the main text or the supplementary materials

