## Supplemental figures for "Reduction of Spermine Synthase Suppresses Tau Accumulation Through Autophagy Modulation in Tauopathy"

**Supplementary figures**

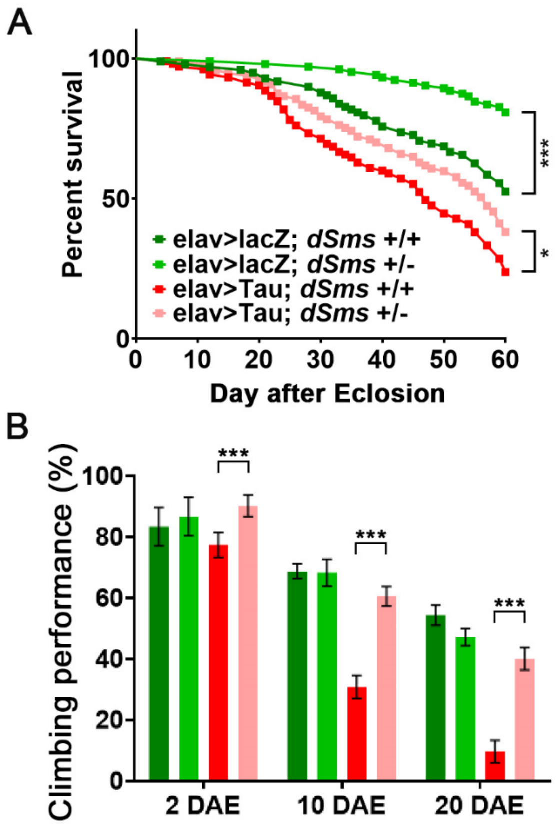


**Figure S1. SMS reduction extends lifespan and improves climbing performance in male Tauopathy flies.** (A) Lifespan of male flies with indicated genotype. n = 99, 104, 105, 97; Log-rank (Mantel-Cox) test. (B) Climbing performance of male flies with indicated genotype at indicated ages. n = 100, 100, 100, 100; two-way ANOVA multiple comparisons. Data represent mean ± SEM.

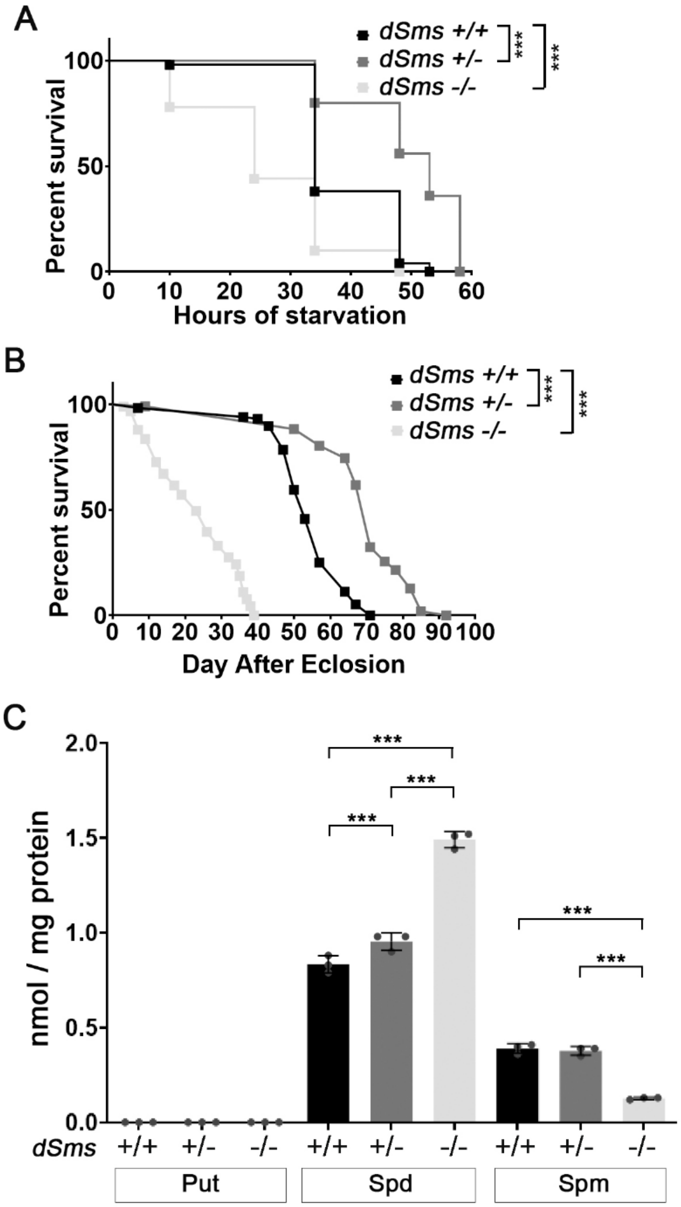


**Figure S2. SMS regulates autophagy function in male flies.** (A) Survival curve of 10 DAE male flies with indicated genotype under starvation. n = 50, 50, 50; Log-rank (Mantel-Cox) test. (B) Lifespan of male flies with indicated genotype. n = 116, 102, 91; Log-rank (Mantel-Cox) test. (C) Polyamine levels of 10 DAE male flies with indicated genotype. Each dot indicates a sample of homogenized mixture of 10 flies. n = 3; two-way ANOVA multiple comparisons. Data represent mean ± SEM. The measurement showed here was done together with that showed in our previous publication [1]. The data of the control and *dSms* -/- flies are shared in these two studies.
